# Human development and inequality shape global threat patterns for terrestrial biodiversity

**DOI:** 10.64898/2026.08.07.743246

**Authors:** Katherine Pulido-Chadid, Adrienne Etard, Antonella Gorosábel, Martin Jung, Louise M. J. O’Connor, Carsten Rahbek, Jonas Geldmann

## Abstract

Biodiversity loss is driven by unsustainable human activities, yet the contextual conditions and underlying drivers of threats remain poorly understood. We assessed how protected areas, socioeconomic conditions, and biophysical factors explain global patterns of threat probabilities across six major threat types and four vertebrate taxa. We identified key explanatory variables and their associations with threats using Extreme Gradient Boosting (XGBoost) and SHapley Additive exPlanations (SHAP). Socioeconomic conditions, specifically human development and income inequality, were the strongest predictors. Their associations were complex and non-linear: notably, high human development index (HDI) was associated with both higher and lower threat probabilities, depending on inequality and regional context. Second, land cover and biophysical variables, such as shrubland cover, tree cover, and elevation range, explained additional, but taxon-specific variation. Finally, protected areas showed limited ability to explain threat patterns. By linking threat probabilities to their contextual and socioecological conditions, we aim to build a better understanding of the systemic drivers of biodiversity loss.

## 1 Introduction

Biodiversity loss has been accelerating for decades because of unsustainable human activities (Barnosky et al., 2011; Ceballos and Ehrlich, 2023; Di Marco et al., 2018; Pimm et al., 2014; Tilman et al., 2017). Since the 1950s, economic development has intensified pressures on nature, compromising ecosystem functioning and resilience in many places (Muys, 2013; Steffen et al., 2015). This has resulted in increased pressure on biodiversity, which in turn jeopardizes nature’s contributions to people (Díaz et al., 2019; Hill et al., 2021; IPBES, 2019). Globally, the main threats to biodiversity are changes in land and sea use, overexploitation of organisms, climate change, pollution, and invasive alien species (IPBES, 2019). Major threats to biodiversity are underpinned by indirect drivers, such as models of production and consumption and technological development, which are all shaped by the complex ways people interact with nature (IPBES, 2025, 2019).

The potential impact of threats varies with the taxonomic group, species ecological characteristics, habitat, landscape features (Bellard et al., 2022; Foden et al., 2013), and socio-environmental contexts (Geldmann, 2023; IPBES, 2019; Maxwell et al., 2020; Schulze et al., 2018). These systemic elements are intrinsically linked to broader socioeconomic conditions, including levels of development, human well-being and equality (Carmenta et al., 2025; Dasgupta & Levin, 2023; Gatiso et al., 2022; Geldmann et al., 2014).

Threat maps have been developed to identify where and to what extent biodiversity is at risk from human activities (Pla et al., 2024; Ridley et al., 2024; Tulloch et al., 2015). Such maps describe the distribution, intensity, or frequency of threats across an area and can be used to identify priority sites for conservation (Brooks et al., 2006; Geldmann et al., 2014; Tulloch et al., 2015). However, threat maps have traditionally been developed using data on human drivers of threats and, thus, often lack information about the impact on species (Balmford et al. 2009). Further, these maps often omit threats for which data at macroecological scales are lacking, such as overexploitation, pollution, and invasive species (Harfoot et al., 2021). To address these limitations, the International Union for Conservation of Nature (IUCN) Red List of Threatened Species provides a powerful data source, as, in addition to evaluating the extinction risk of assessed species, it also identifies the threats affecting each species (IUCN, 2025). However, threat evaluations are done at the species level, and the IUCN Red List does not specify the precise location where these threats occur within a species’ range. To address this limitation, Harfoot et al. (2021) estimated the probability of impact of a given threat in an area by looking at the proportion of all species in that area affected by a threat, while accounting for species’ range sizes to ensure that species with smaller ranges, and therefore larger spatial certainty of the spatial location of a threat, were given more weight (Harfoot et al. 2021). This approach enabled global, spatially explicit assessments of how different threats affect vertebrate groups (Farooq et al., 2024; Harfoot et al., 2021).

Protected areas (PAs) are one of the most important tools in nature conservation; when effective, they can help to reduce threats to biodiversity (Schulze et al., 2018). However, PAs are often designated in remote places or regions of low economic value, where accessibility is limited, and threats are comparably low (Joppa & Pfaff, 2009, 2011). This spatial bias can overestimate PA effectiveness, as some observed conservation outcomes may reflect inherent baseline characteristics or a lack of disturbance, rather than the impact of protection itself (Cazalis et al., 2020; Geldmann et al., 2025; O’Garra et al., 2025). Meanwhile, threats to biodiversity continue to intensify both inside and outside PA boundaries (Geldmann et al., 2014; J Geldmann et al., 2019; Jones et al., 2018; Pulido-Chadid et al., 2023; Schulze et al., 2018). PA effectiveness also depends on the surrounding context, institutional capacity and management enforcement (Durán et al., 2022; Geldmann et al., 2025; Graham et al., 2021; Wauchope et al., 2022, Pulido-Chadid et al., 2025). Understanding not only where threats occur but also the broader socio-environmental conditions in which they operate is therefore essential.

Previous work has shown that the probability of threat impact is not aligned with PA coverage (Pulido-Chadid et al., 2025), highlighting potential mismatches between the placement of PA and areas where species face the highest threats (Negret et al., 2024). However, little is known about the contextual factors that influence these outcomes. Although it is well established that threats are shaped by ecological, landscape, socioeconomic and governance characteristics (IPBES, 2025, 2019), we still have a limited understanding of how these factors influence the spatial distributions of threats. In particular, how these relationships vary across threat types, taxonomic groups, and regional development contexts remains poorly understood.

To address this gap, analytical tools capable of modelling complex and nonlinear relationships are needed. Machine learning (ML) techniques are suitable for this purpose, as they can be trained directly on the data without meeting explicit assumptions about the underlying data-generating process (Cutler et al., 2007; Elith et al., 2008). Another advantage of ML is the ability to uncover patterns, relationships and interactions between predictors and response variables (Elith et al., 2008; Pichler and Hartig, 2023; Yu et al., 2025). Here, we employ XGBoost, a high-performance tree-based regression algorithm, together with the ML technique SHAP to identify the top predictors explaining threat patterns and their influence across taxa and regions. We focus on 14 covariates, including socioeconomic conditions, biophysical variables and PA coverage, and assess their associations with six major threats to terrestrial vertebrates: agriculture, hunting and trapping, logging, invasive species and diseases, pollution, and urbanization. Our analysis builds on global threat probability maps developed by Harfoot et al. (2021) and Farooq et al. (2024), derived from the International Union for Conservation of Nature (IUCN) distribution data for 33,379 species of amphibians, birds, mammals, and reptiles. Our objectives are to (1) identify key variables explaining threat patterns across taxa, (2) assess differences and similarities among threats and taxa, and (3) examine the direction and magnitude of the top predictors across regions.

We expect a strong influence of socioeconomic conditions and land-use type on threat probabilities (Cincotta et al., 2000; IPBES, 2019), with variations between taxonomic groups and threat types. Previous studies have shown little relationship between PA coverage and threats in general (Pulido-Chadid et al., 2025), and here we test whether this is also verified for threat patterns across regions and taxa. By linking threat probabilities to their geographical and socioeconomic context, as well as current levels of protection, we aim to provide new insights into the underlying drivers of threats to biodiversity and the role of PAs in shaping these patterns.

## 2 Methods

### 2.1 Data and preprocessing

#### 2.1.1 Threat maps

We used the threat probability maps developed by Harfoot et al. (2021) and Farooq et al. (2024). These threat maps were based on amphibian, mammal, and reptile species ranges from IUCN Red List version 2022-1, and bird species distribution from BirdLife International (version 2021.1). The maps estimate the probability that a species occurring in a grid cell is impacted by a particular threat. To do so, species range maps were combined with IUCN Red List threat assessments in a probabilistic framework that accounts for the fact that threats are recorded at the species level rather than spatially within species ranges, thereby inferring the most likely spatial distribution of threat impacts from patterns across thousands of species. These data comprised 33,379 terrestrial species, covering 7,073 amphibians, 10,959 birds, 5,581 mammals, and 9,766 reptiles. The six major threats investigated here were: 1) agriculture, 2) hunting and trapping (hereafter “hunting”), 3) invasive and other problematic species, genes & diseases (hereafter “invasive species”), 4) logging, 5) pollution, and 6) urbanization for each of the four taxa. In total, this resulted in 24 individual threat maps, quantifying the probability of impact of a specific threat for a specific taxon. These data used an equal-area Mollweide projection in a 50 x 50 km spatial resolution. Following Harfoot et al. (2021), only cells with more than ten species were considered in the analysis to reduce uncertainty in areas with a low number of species. As a result, the analysis included a sample size of 50,624 grid cells for mammals, 54,979 for birds, 36,252 for reptiles, and 22,936 for amphibians.

### 2.2 Explanatory variables

To assess the importance of different predictors in explaining threat patterns to terrestrial vertebrates, we compiled a set of spatial covariates covering Land Use and Land Cover (LULC), elevation, socioeconomic indicators, and PA coverage (Table 1). These variables were selected because they capture underlying drivers of threats, anthropogenic pressures, habitat characteristics, and ecological dynamics (Mouillot et al., 2024; Theobald et al., 2025).

**Table 1.**
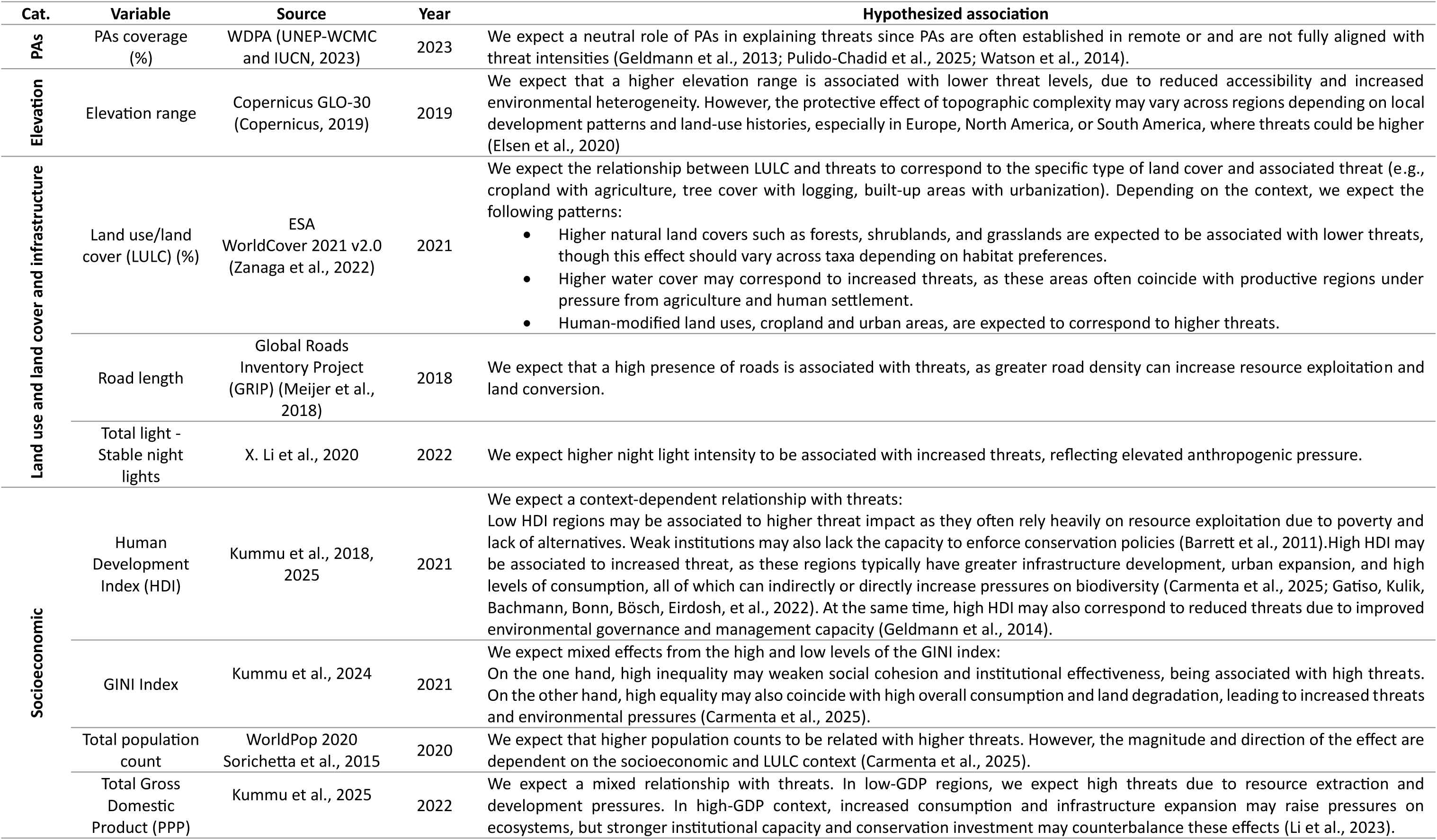
Explanatory variables for modeling spatial patterns of threats to terrestrial vertebrates.

| Cat. | Variable | Source | Year | Hypothesized association |
| --- | --- | --- | --- | --- |
| PAs | PAs coverage (%) | WDPA (UNEP-WCMC and IUCN, 2023) | 2023 | We expect a neutral role of PAs in explaining threats since PAs are often established in remote or and are not fully aligned with threat intensities (Geldmann et al., 2013; Pulido-Chadid et al., 2025; Watson et al., 2014). |
| Elevation | Elevation range | Copernicus GLO-30 (Copernicus, 2019) | 2019 | We expect that a higher elevation range is associated with lower threat levels, due to reduced accessibility and increased environmental heterogeneity. However, the protective effect of topographic complexity may vary across regions depending on local development patterns and land-use histories, especially in Europe, North America, or South America, where threats could be higher (Elsen et al., 2020) |
| Land use and land cover and infrastructure | Land use/land cover (LULC) (%) | ESA WorldCover 2021 v2.0 (Zanaga et al., 2022) | 2021 | We expect the relationship between LULC and threats to correspond to the specific type of land cover and associated threat (e.g., cropland with agriculture, tree cover with logging, built-up areas with urbanization). Depending on the context, we expect the following patterns:<br>Higher natural land covers such as forests, shrublands, and grasslands are expected to be associated with lower threats, though this effect should vary across taxa depending on habitat preferences.<br>Higher water cover may correspond to increased threats, as these areas often coincide with productive regions under pressure from agriculture and human settlement.<br>Human-modified land uses, cropland and urban areas, are expected to correspond to higher threats. |
|  | Road length | Global Roads Inventory Project (GRIP) (Meijer et al., 2018) | 2018 | We expect that a high presence of roads is associated with threats, as greater road density can increase resource exploitation and land conversion. |
|  | Total light - Stable night lights | X. Li et al., 2020 | 2022 | We expect higher night light intensity to be associated with increased threats, reflecting elevated anthropogenic pressure. |
| Socioeconomic | Human Development Index (HDI) | Kummu et al., 2018, 2025 | 2021 | We expect a context-dependent relationship with threats:<br>Low HDI regions may be associated to higher threat impact as they often rely heavily on resource exploitation due to poverty and lack of alternatives. Weak institutions may also lack the capacity to enforce conservation policies (Barrett et al., 2011). High HDI may be associated to increased threat, as these regions typically have greater infrastructure development, urban expansion, and high levels of consumption, all of which can indirectly or directly increase pressures on biodiversity (Carmenta et al., 2025; Gatiso, Kulik, Bachmann, Bonn, Bösch, Eirdosh, et al., 2022). At the same time, high HDI may also correspond to reduced threats due to improved environmental governance and management capacity (Geldmann et al., 2014). |
|  | GINI Index | Kummu et al., 2024 | 2021 | We expect mixed effects from the high and low levels of the GINI index:<br>On the one hand, high inequality may weaken social cohesion and institutional effectiveness, being associated with high threats. On the other hand, high equality may also coincide with high overall consumption and land degradation, leading to increased threats and environmental pressures (Carmenta et al., 2025). |
|  | Total population count | WorldPop 2020 Sorichetta et al., 2015 | 2020 | We expect that higher population counts to be related with higher threats. However, the magnitude and direction of the effect are dependent on the socioeconomic and LULC context (Carmenta et al., 2025). |
|  | Total Gross Domestic Product (PPP) | Kummu et al., 2025 | 2022 | We expect a mixed relationship with threats. In low-GDP regions, we expect high threats due to resource extraction and development pressures. In high-GDP context, increased consumption and infrastructure expansion may raise pressures on ecosystems, but stronger institutional capacity and conservation investment may counterbalance these effects (Li et al., 2023). |

All layers were harmonized to a global 50 × 50 km grid in the Mollweide projection, aligned to the spatial extent and resolution of the threat probability layers. Since the threat maps included species information up to 2022, we used data from 2022 where available, or the closest year otherwise (Table 1). Data on human population count, LULC, nighttime lights, and elevation were processed in Google Earth Engine (GEE), and the remaining variables were processed in R v4.2.3 (R Core Team, 2021) using the sf (Pebesma, 2018; Pebesma and Bivand, 2023), raster (Hijmans, 2023), terra (Hijmans, 2024) and exactextractr (Baston, 2023) packages.

#### 2.2.1 Protected areas cover

We calculated proportional PA coverage for each grid cell using terrestrial and coastal polygons from the September 2023 release of the World Database on Protected Areas (WDPA; UNEP-WCMC and IUCN, 2023). Due to the unavailability of public WDPA data for China, India, and Turkey, we excluded them from the analysis. We retained sites that met the PA definition: “designated” (legally established), “established” (effective by other means), or “not reported” (UNEP-WCMC, 2019). To ensure the analysis focused on terrestrial PAs, we intersected these polygons with a global landmass mask from OpenStreetMap (2019) to exclude marine areas. In total, 255,848 sites were rasterized into the 50 km Mollweide grid. For each grid cell, we then calculated the fraction of the cell covered by PAs considering: (i) total PA coverage, (ii) strict PA coverage (IUCN categories I–II), and (iii) non-strict PA coverage (all other categories).

#### 2.2.2 Elevation range

Elevation range captures topographic complexity, which can influence both habitat heterogeneity and human accessibility (Elsen and Tingley, 2015; Joppa and Pfaff, 2009; Stein et al., 2014). Elevation data was obtained from Copernicus GLO-30 Digital Elevation Model (DEM) given in 30 m resolution (Copernicus, 2019). In GEE, we identified the maximum and minimum elevation in each 50 x 50 km cell. We then calculated the elevation range as the difference between them.

#### 2.2.3 Land use and land cover percentage

We derived the percentage covered by LULC types from ESA WorldCover 2021 v2.0 (Zanaga et al., 2022) within each 50 x 50 km grid cell. Using GEE, we counted the 10 × 10 m pixels within each grid cell to calculate the percentage of each LULC type. All eleven types were retained: tree cover, shrubland, grassland, cropland, built-up areas, bare vegetation, permanent water bodies, mangroves, moss/lichen, snow/ice, and herbaceous wetlands. Mangroves, moss/lichen, snow/ice, and herbaceous wetlands were grouped as “other land uses,” which together accounted for ∼5% of the dataset.

#### 2.2.4 Road length

The presence of roads facilitates human access, habitat fragmentation, and ecosystem degradation (Ibisch et al., 2016). We used the Global Roads Inventory Project (GRIP) road-density raster (GLOBIO, 2018; Meijer et al., 2018) to compute the total road length (km) within each 50 × 50 km grid cell. The GRIP product reports the combined road densities of highways, primary, secondary, tertiary, and local roads per unit area (m/km²) at a native 5 arcminute resolution in WGS 84. We reprojected the road density layer into an equal-area Mollweide projection, calculated each pixel’s area, and multiplied it by the density to estimate road length in km per pixel. Then, we overlaid the 50 × 50 km grid and summed road lengths within each cell.

#### 2.2.5 Nighttime lights

The nighttime light illustrates the intensity of human emitted light and can serve as a proxy for human activity, habitat fragmentation, and infrastructure development (Kong et al., 2022; G. Li et al., 2020). To estimate the cumulative and mean radiance value for each cell, we used the annual composite from the Visible Infrared Imaging Radiometer Suite (VIIRS) Day/Night Band (DNB), available in GEE (Elvidge et al., 2021). This product provides pre-processed, cloud-masked radiance at 500 m resolution in units of nanoWatts per square centimeter per steradian (nW cm⁻² sr⁻¹). We filtered the image collection to the 2022 period (2022-01-01 to 2023-01-01) and selected the average masked band, which represents the mean radiance after removing cloud-contaminated and other low-quality observations. We calculated the total radiance for each grid cell.

#### 2.2.6 Human population count

Human population indicates demographic pressure and demand for land, resources, and infrastructure (Cincotta et al., 2000; Theobald et al., 2025; Venter et al., 2016). To calculate the total number of inhabitants within each grid cell, we used the WorldPop 2020 population dataset (Sorichetta et al., 2015; WorldPop, 2020) and aggregated all pixel values with a sum within each 50 × 50 km grid cell.

#### 2.2.7 Total Gross Domestic Product (GDP)

Gross Domestic Product (GDP) represents the total value of final goods and services produced within a region, reflecting income and expenditure (Callen, 2012; Kummu et al., 2018). GDP is often associated with resource use and infrastructure expansion (Li et al., 2023). Kummu et al. (2025) developed a globally harmonized dataset of GDP per capita (PPP) at a 30-arc-minute resolution from 1990 to 2022. This dataset was downscaled to administrative level 2 (i.e., municipality), considering 43,501 units. By combining this downscaled product with population count data, they estimated total GDP (PPP) for each cell on an annual basis. We extracted the 2022 layer and reprojected it to the 50 × 50 km Mollweide grid using bilinear resampling to obtain the total GDP (PPP) within each 50 × 50 km grid cell.

#### 2.2.8 Human Development Index

The Human Development Index (HDI) summarizes average achievement across three dimensions: life expectancy, education (mean years of schooling), and standard of living measured by gross national income per capita (UNDP, 2025). HDI is a measure of welfare and development (Chen et al., 2023), and influences governance, land use, and conservation capacity (Gatiso, et al., 2022; Geldmann et al., 2019; Naidoo & Adamowicz, 2001). We used the global HDI time-series dataset from Kummu et al. (2018), which provides annual HDI at 5′-arc-minute resolution for 1990–2021. We extracted and resampled the 2021 layer using bilinear interpolation and computed the modal HDI value within each grid cell. Given the coarse resolution and smooth spatial variation of the original dataset, we found that median and mode aggregation produced values almost identical to the mode (Pearson’s r = 0.999), indicating that the choice of aggregation statistic has a negligible influence on the results.

#### 2.2.9 GINI index

The GINI index measures the extent to which the distribution of income or consumption among households within an economy deviates from an equal distribution. Measured from 0 to 100, a GINI index of 100 implies perfect inequality, where a single individual holds all the income (The World Bank Group, 2025). Income inequality serves as a proxy for broader societal conditions, including social justice, political stability, democratic governance, and corruption (Chrisendo et al., 2024; Ferreira et al., 2022). Thus, it can also affect conservation outcomes by shaping access to resources, enforcement of environmental laws, and the capacity for collective action for conservation (Holland et al., 2009; Kashwan, 2017). We used the global subnational GINI index dataset from Chrisendo et al. (2024), which provides an annual GINI index at a five arc-minute resolution from 1990 to 2021. We extracted the 2021 layer and reprojected using bilinear resampling. We then applied a modal extraction to assign each grid cell the most frequent GINI value.

#### 2.2.10 Variable screening

To ensure the independence and relevance of predictors, we conducted a two-step preliminary analysis using the R package spatialRF (Blas, 2021). First, we assessed Pearson pairwise correlations among variables and removed those with correlation coefficients above 0.7 (Dormann et al., 2013). Then, we further assessed multicollinearity using the Variance Inflation Factor (VIF), excluding variables with VIF values greater than 5. While high correlation indicates a strong relationship between two variables, multicollinearity occurs when two or more predictors are linearly related (Dormann et al., 2013). We screened the candidate variables once, independently of threat and taxon, which resulted in the removal of GDP and percentage of built-up areas due to high pairwise correlation with other variables, and percentage of bare vegetation due to high multicollinearity and likely redundancy with other land-cover predictors (Dormann et al., 2013; O’Brien, 2007). The final selection included 14 variables: HDI, GINI, total population, roads length, total light, elevation range, and percentage covered by: PAs or strict-PAs and non-strict PAs, water, cropland, grassland, shrubland, tree cover, and other LULCs. This variable set was used across all taxa to ensure comparability.

### 2.3 Explaining threat patterns using XGBoost and SHAP

#### 2.3.1 Extreme Gradient Boosting and Shapley Additive Explanations

We used XGBoost, a gradient-boosted decision tree algorithm, to perform regression analysis and identify the variables most strongly associated with threat probabilities (Chen and Guestrin, 2016). Gradient-boosted trees handle mixed predictor types and missing values, require minimal preprocessing, and capture complex nonlinear relationships, making them well-suited for ecological data (Friedman, 2001; Jung, 2023; Manley et al., 2023). A single decision tree partitions the data with a sequence of if-then rules, whereas boosting aggregates many shallow trees into an ensemble that is both more accurate and less prone to over-fitting (Elith and Leathwick, 2009). XGBoost is known for its high predictive performance, speed, and flexibility, and is widely used in machine learning competitions and real-world applications (Chen and Guestrin, 2016). Here, we used XGBoost in a regression framework to identify which predictors best explained variation in threat probabilities.

To interpret the XGBoost models, we applied SHAP (Shapley Additive Explanations), a post-hoc explainability method based on cooperative game theory (Lundberg and Lee, 2017). SHAP assigns each predictor a contribution value for every individual prediction (local explanations). Aggregating these across all predictions yields the global variable importance summarized as the mean absolute value (mean |SHAP|). This allows for assessing both the direction and magnitude of predictor contributions while accounting for multicollinearity and interactions (Lundberg et al., 2020; Lundberg and Lee, 2017; Zhang et al., 2023).

#### 2.3.2 Base model configuration

We used XGBoost to model the probability of threat impact (0–1) in each 50 × 50 km grid cell as a function of the 14 selected independent variables. We modeled six threats independently: agriculture, hunting, logging, pollution, invasive species, and urbanization, as well as the cumulative threats index (the sum of all six threats) for amphibians, birds, mammals, and reptiles. In all cases, the probability of threat impact was the response variable. We aimed to identify which variables most strongly explained spatial variation in threat probabilities and the degree to which the predictors contributed to the modeled threats. To achieve this, we applied SHAP, which quantifies the contribution of each predictor to individual model predictions.

##### Model training and performance

Each model was fitted to one of seven outcomes per taxon: the six individual threats and the cumulative threat, defined as the sum of the six individual threat probabilities within each grid cell. Data were split into 80% training and 20% testing sets. We used a grid search to identify the best-performing model for each outcome, in which the model is trained and evaluated on all combinations of selected hyperparameters to find the settings that give the best performance. The hyperparameters we tested included the learning rate (which controls how quickly the model learns from the training data), maximum tree depth (which determines model complexity), number of boosting rounds, and subsampling rates for both training instances and predictor columns (the fraction of observations to be randomly sampled for each tree) (xgboost developers, 2025). The objective function of each trained model aimed to minimize the root mean squared error (RMSE); a measure of how far predictions are from observed values on average. We applied early stopping to avoid overfitting, halting training if the RMSE did not improve for 10 consecutive rounds. Once the best settings were identified, we evaluated each model on the test set using RMSE and the coefficient of determination (R²).

##### Assessing variable contributions: importance and direction

We used SHAP to evaluate the role of each predictor in the models. SHAP values were computed from the final trained model and applied to the full dataset, providing local explanations of how each variable contributed to the predicted threat probability in every 50 × 50 km grid cell. We distinguished between two complementary aspects captured by SHAP values: First, the magnitude of contributions (variable importance), i.e. how each predictor contributed to the predicted threat probability in each grid cell. To quantify global variable importance, we calculated mean |SHAP|. Larger mean |SHAP| values indicated a stronger overall contribution to the model’s predictions. “Most important” or “top” predictors refer to those variables with the highest mean |SHAP| values. Second, we considered the direction of contributions. Signed SHAP values indicated whether a predictor was associated with higher or lower predicted threat probability.

Analyses were conducted in R using xgboost (Chen et al., 2025), SHAPforxgboost (Liu and Just, 2023), tidyverse (Wickham et al., 2019) and sf packages.

#### 2.3.3 Analytical framework

We applied the same base configuration to two sets of models with different targets and objectives. First, to identify the most important predictors for each threat–taxon combination, we fitted 24 independent models (6 threats × 4 taxa). Within each model, we ranked predictors by mean ∣SHAP∣ and treated a predictor as important when it ranked among the top three, then applied a threshold of mean ∣SHAP∣ > 0.025, selected based on a natural break in the distribution of values across taxa to ensure comparability (Figure S2). Second, to evaluate how predictors contributed to cumulative threats within each taxon, we used SHAP summary plots to examine both the importance and the direction of predictors. Additionally, we mapped the predictor with the largest local SHAP value on each grid cell, retaining its sign to investigate its association with threat levels. To further explore top predictors, we generated SHAP dependence plots for the most important variables. Because HDI and GINI showed complex, non-monotonic associations with cumulative threats, we used interaction plots to examine how GINI’s effect varied across the HDI gradient. To aid interpretation, predictors were grouped into four categories: (1) Socioeconomic: HDI, GINI, total population; (2) LULC: percentage covered by water, cropland, grassland, shrubland, tree cover, other LULCs, roads length, and total light; (3) Elevation range, and (4) PA coverage.

### 2.4 Model performance

The models generally achieved strong predictive performance across taxa and threat types, for both individual and cumulative threats. R² values ranged from 0.66 to 0.94, indicating that the models explained between 66% and 94% of the variance in observed threat probabilities. RMSE values ranged from 0.02 to 0.11 for individual threats and from 0.11 to 0.35 for cumulative threats, indicating that predictions were close to the observed values (Table S1).

## 3 Results

### 3.1 Predictors of threats

Socioeconomic factors, particularly HDI and GINI, emerged as the most important variables explaining threat patterns across all threats and taxa. Of the 14 predictors evaluated, HDI ranked among the top three in all 24 threat–taxon models and GINI in 80% of them, followed by elevation and LULC categories whose relevance was taxon-specific (Figure 1).

**Figure 1.**
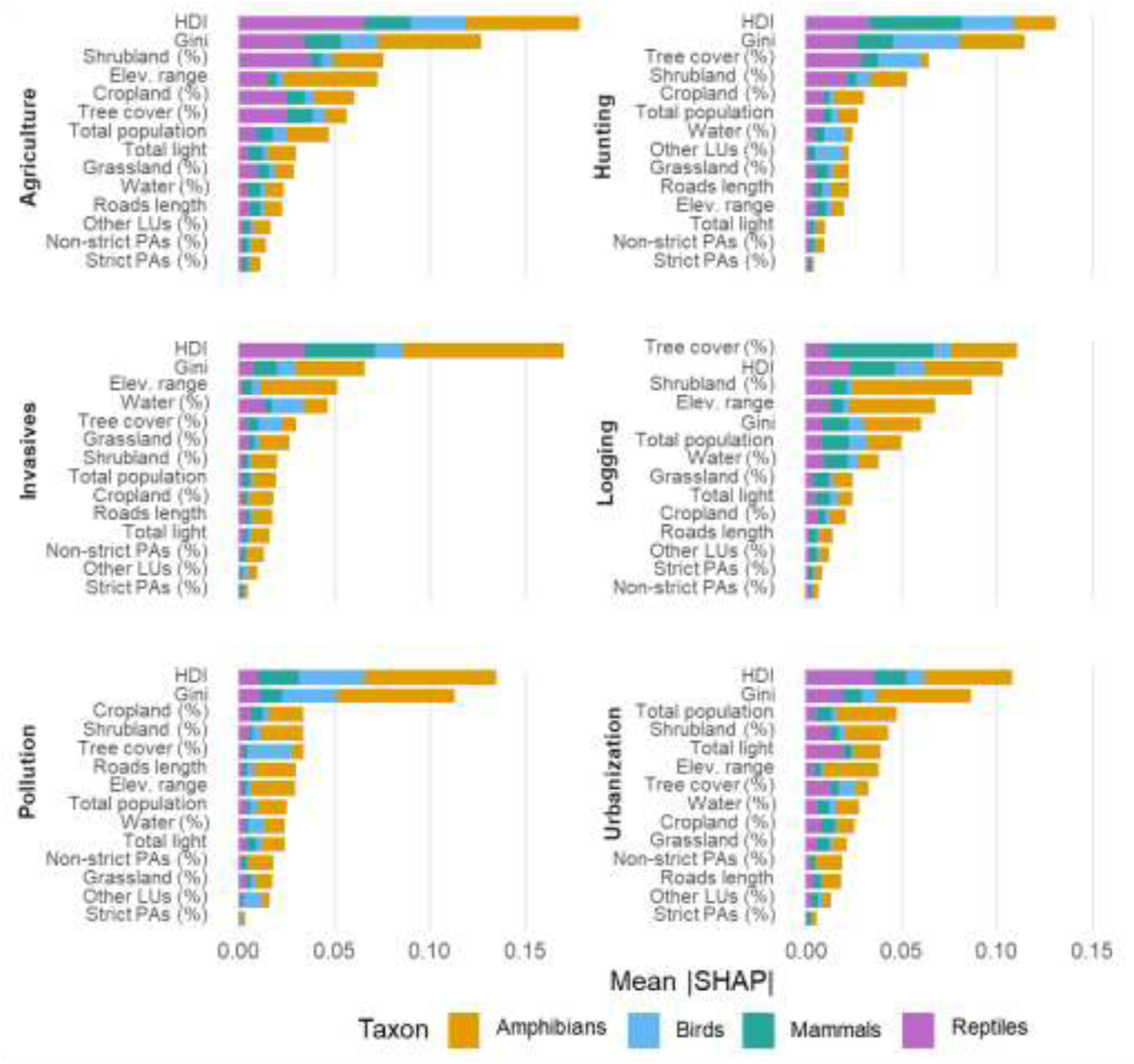
Variable importance of predictors across threats and taxa. Predictors are ranked by mean |SHAP|, which summarizes the average contribution of each variable to model predictions across all grid cells. Panels correspond to threats and colors indicate taxa, so each colored segment is the mean |SHAP| of one threat–taxon model (e.g., agriculture– amphibians, logging–mammals). Segments are stacked by taxon for comparison and are not summed. Results are based on individual threat-taxon, summarizing 24 models.

HDI was among the top predictors explaining all threats to amphibians except from hunting, as well as agriculture and hunting to reptiles, mammals, and birds; invasive species to reptiles and mammals; pollution to birds; and urbanization to reptiles. GINI was important for amphibians across all threats, and in hunting for birds and reptiles, in pollution for birds, and agriculture for reptiles (Figure 1).

LULC and elevation showed taxon-specific importance. Shrubland cover was important for the threat posed by agriculture for amphibians and reptiles, and by logging for amphibians. Tree cover was important for amphibians and mammals in logging, and for reptiles in hunting. Some predictors were especially taxon-specific. For example, cropland cover was important for predicting the probability of impact from agriculture for reptiles, and human population count in urbanization for amphibians. Elevation range was a top predictor for most threats to amphibians (Figure 1).

PA coverage, both strict and non-strict, had low importance across all models (Mean ∣SHAP∣ < 0.01), indicating a limited role of PA coverage in explaining predicted threat probabilities (Figure 1).

### 3.2 Predictors importance and direction on cumulative threats

Predictor contributions to cumulative threats varied across taxa both in magnitude and direction (Figure 2). HDI was consistently a top predictor, but its effects were non-linear and context dependent: high HDI values were associated with both high and low cumulative threats (Figure 2), with dependence plots confirming these patterns (Figure S8). This context dependency was largely driven by interactions with inequality (GINI), which amplified threats in high-HDI regions but had weak or even negative effects in lower-HDI areas (Figure S9). GINI showed an overall negative association, with lower inequality associated with higher predicted threats, particularly for amphibians and mammals (Figure 3 and Figure S8).

**Figure 2.**
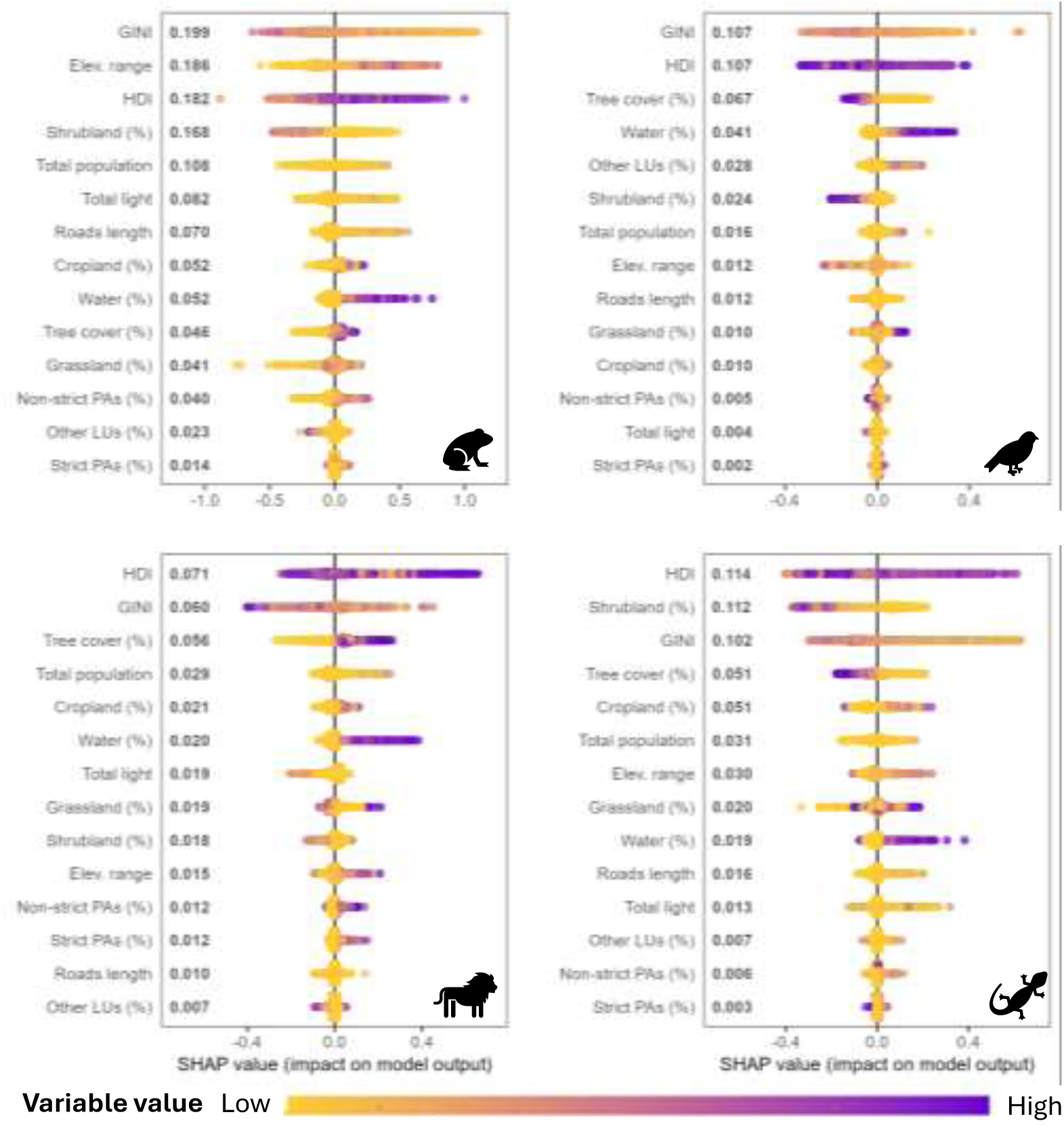
SHAP summary plots for cumulative threat models for amphibians, birds, mammals, and reptiles. Predictors are ranked by mean ∣SHAP∣ values, showing their overall contribution to the model in descending order. The horizontal position of each point shows its SHAP value in a grid cell, indicating whether lower or higher predictor values were associated with an increase (positive SHAP) or decrease (negative SHAP) in predicted cumulative threat probability. Point colors represent the actual predictor values (yellow = low, purple = high). The horizontal spread of points reflects the variability of predictor effects across grid cells, with wider spreads indicating stronger and more heterogeneous impacts on predictions

**Figure 3.**
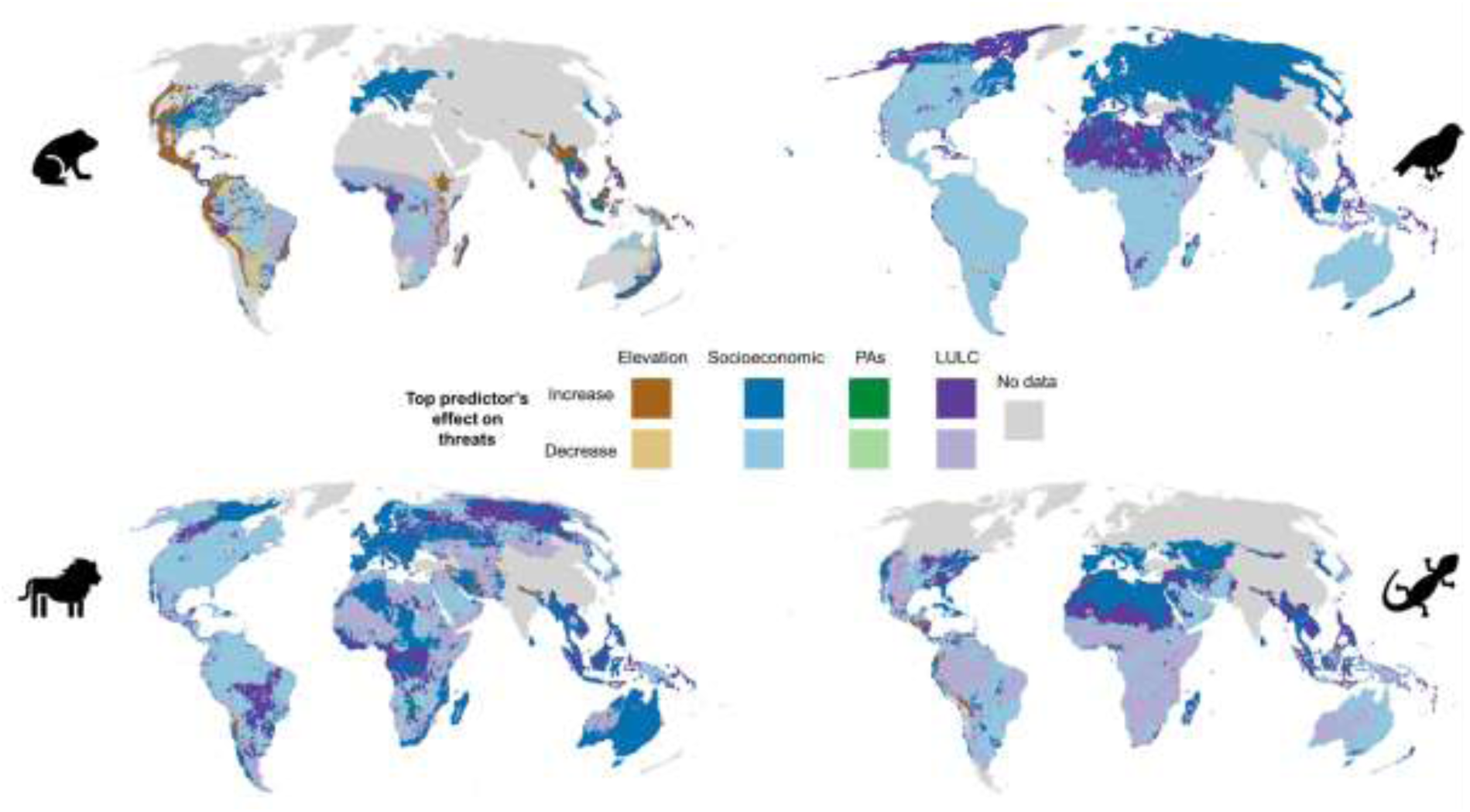
Magnitude and direction of top predictors associated with predicted cumulative threats. The maps show, for each grid cell, the predictor with the largest SHAP contribution, retaining its sign (positive or negative) to indicate whether the predictor was associated with higher or lower predicted threat probability. Positive SHAP values indicate an association with increased predicted threats, while negative SHAP values indicate an association with reduced predicted threats. Grey areas indicate no data.

For LULC variables, some consistent patterns emerged: greater shrubland cover was generally associated with lower threats, while higher water cover was linked to high threats across taxa (Figure 2). Other land cover variables showed more taxon-specific responses. For instance, greater tree cover was associated with lower threats for birds and reptiles but with increased threats for mammals (Figure 2). Human population count and nighttime light showed mixed contributions, with both positive and negative effects. Most of this variability was associated with low threat probability values, particularly for amphibians and reptiles (Figure 2).

### 3.3 Spatial patterns of the top predictors of cumulative threats

Socioeconomic indicators were the most widespread top predictors across taxa for cumulative threats, ranking as the top predictor in 47% of the mapped area for amphibians, 79% for birds, 61% for mammals, and 51% reptiles (Figure 3). Their associations varied geographically: In Europe and Asia, characterized by high human development and low to medium inequality, socioeconomic variables were linked to higher threat, whereas in Central Africa and South America, where development and inequality both span wide ranges, they were associated with lower threats for amphibians, birds, and reptiles, and mixed responses for mammals (Figure 3 and Figure S1-m).

LULC variables were the second most frequent top predictors, accounting for 30% of the mapped area for amphibians, 38% for mammals, 21% for birds and 48% for reptiles. Shrubland cover was the most frequent LULC type associated with lower threats for amphibians (17%) and reptiles (24%), particularly in regions such as the Caatinga and Patagonian steppe (South America), sub-Saharan Africa, and western Australia, where shrubland cover was high (Figure 3 and Figure S1-d). Tree cover showed opposing patterns across taxa. For mammals, it was associated with higher threats in regions of extensive cover such as Eastern Europe and Central Africa, whereas for reptiles high tree cover corresponded to lower threats, as in the Amazon and Congo basin (Figure 3 and Figure S1-a).

For amphibians, a higher elevation range was associated with higher threats across 14.2% of the mapped area, concentrated in montane regions including the Andes, the Himalayas, parts of Central America, the Ethiopian Highlands, and Southeast Asia (Figure 3 and Figure S1-i).

PAs were rarely the top predictor (less than 0.5% of grid cells), with inconsistent associations: PA cover was linked with lower threats for amphibians in North America but higher threats for mammals in parts of Southern Africa and Europe (Figure 3). The direction and magnitude of each predictor are detailed in Figure S10.

## 4 Discussion

Socioeconomic factors, especially human development and income inequality, were the variables most strongly associated with threat patterns to terrestrial vertebrates, ranking as the top predictor in over half of the mapped area. These correspond to the indirect drivers of biodiversity loss defined by IPBES (2019) and our findings align with previous research linking socioeconomic context to both threats and conservation responses. For example, Brabant et al. (2025) identified HDI as the most relevant predictor of mining-related threats to freshwater species, while Mouillot et al. (2024) found that PAs were disproportionately represented in areas with higher HDI and greater NGO presence.

Although the relationship between socioeconomic context and biodiversity loss has been widely covered in the literature, our findings suggest that threats arise in a complex and context-dependent interplay among economic growth, human development, and inequality. This complexity is also recognized in previous studies, which have found contrasting relationships between socioeconomic drivers and threats to biodiversity. For example, York et al. (2003) identified the overall size of an economy as the primary driver of environmental impacts, while Cafaro et al. (2022) argued that poverty and population growth can be contributors to biodiversity loss (but see response by Green et al. (2022)). Conversely, studies have also found positive associations between socioeconomic indicators and threats to biodiversity: Naidoo & Adamowicz (2001) reported that the number of threatened species increased with higher Gross National Product per capita, while Mikkelson et al. (2007) observed the same pattern with income inequality (GINI).

Our results indicate no single linear relationship between human presence and threats. Low human population and nighttime light were linked to both lower and higher threat probabilities, highlighting a context-dependent relationship where socioeconomic and land-use factors mediate human impact. This reinforces the idea that threats to biodiversity are not driven by human population density but by complex socioeconomic factors and consumption patterns (Green et al., 2022). Similarly, the associations between threats and development and between threats and inequality were complex. In highly developed regions with low inequality, such as parts of Europe and Asia, threats were often higher, likely reflecting a longer and more intensive land-use history, infrastructure development, and resource consumption (Carmenta et al., 2025; Gatiso et al., 2022a). In contrast, other regions with high human development exhibited lower probability of threat impact, possibly due to stronger governance, management capacity and conservation responses (Gatiso et al., 2022b; Geldmann et al., 2014; Mammides, 2020). These context-dependent associations underline that development and equality are not inherently protective if accompanied by intensive land use, high resource consumption, and long histories of natural resource exploitation (IPBES, 2019; Wiedmann et al., 2020). Our work highlights that the complex interplay between socioeconomic conditions, land use, consumption patterns, and governance likely influences threats.

Biophysical conditions were the next most important variables, ranking as the top predictor in more than one-third of the mapped area, often with taxon-specific patterns. Among these, higher shrubland cover was consistently associated with lower threats for amphibians and reptiles. This aligns with previous studies reporting that greater shrubland cover supports higher species richness and evenness for amphibians and reptiles in arid and semi-arid regions (Owen et al., 2024). This pattern is consistent with the role of habitat loss as a critical threat to amphibians and reptiles. For instance, many threatened amphibian species with small geographical ranges tend to have a lower proportion of suitable habitat remaining (Cordier et al., 2021; Ficetola et al., 2015; Luedtke et al., 2023).

By contrast, tree cover showed divergent associations across taxa, reflecting differences in habitats and species vulnerability to human pressures (Bellard et al., 2022). For reptiles and birds, greater tree cover tended to be associated with lower threats, while for mammals, the opposite pattern emerged, particularly in forested regions of Eastern Europe and Middle Africa. These patterns may be explained by the presence of managed forests or plantations (Jung et al., 2022), or high hunting pressure in intact or semi-intact forests (Benítez-López et al., 2017; Redford, 1992). Large-bodied mammals are especially vulnerable to hunting due to their higher detectability, large range sizes, and slower reproduction rates (Benítez-López et al., 2017; Ripple et al., 2016).

Greater elevation variability was associated with increased threats to amphibians in the Andes, the Himalayas, parts of Central America, the Ethiopian Highlands, and Southeast Asia, which are all under intense human pressure (Elsen et al., 2020). This pattern likely reflects the exposure of amphibians in montane environments, where species tend to have smaller geographic ranges, specialized habitat requirements, and limited dispersal capacity. These characteristics increase their vulnerability to both climatic and land-use changes, making high-elevation regions potential hotspots of extinction risk (Guirguis et al., 2023; Hof et al., 2011).

Our analysis showed that PAs, regardless of strictness, contributed little to explaining threat patterns: PA coverage emerged as a top predictor in less than 0.5% of global land area. This supports previous studies showing that PAs are often located in areas of low economic value, limited accessibility, or low human pressure rather than where protection is most needed (Joppa and Pfaff, 2009; Negret et al., 2024; Pulido-Chadid et al., 2025).

### 4.1 Limitations and uncertainties

While our analysis provides a global view of the socioeconomic and biophysical factors associated with threats to biodiversity, interpreting the results requires careful consideration. The threat maps are based on the IUCN Red List range maps, which are expert-based and generalized, making them susceptible to commission and omission errors and biased toward well-studied regions (Di Marco et al., 2017; Ficetola et al., 2015; Hayward et al., 2015). Variation in the age, completeness, and frequency of IUCN assessments can add further uncertainty, as coverage depends heavily on available resources, with higher-income countries generally better represented (Cazalis et al., 2022). This may partly explain why socioeconomic indicators emerged as strong predictors in our models. Additionally, threat probability maps assume a uniform value across the entire species range, lacking spatial precision on where threats actually occur (Farooq et al., 2024; Harfoot et al., 2021). This is particularly problematic in regions where species range sizes are large and species richness is low, as in the Sahara or the high Arctic, where threats may not be locally relevant but affect species elsewhere in their range (Pulido-Chadid et al., 2025).

The 50 × 50 km resolution of our analysis is appropriate for identifying broad global patterns but not for capturing fine-scale processes or guiding local actions. This coarse scale was chosen to avoid artificial precision given the uncertainty of IUCN range maps (Di Marco et al., 2017; Harfoot et al., 2021; Pulido-Chadid et al., 2025). However, it dilutes spatial heterogeneity, assumes homogeneity within each cell, and mismatches the often small, scattered distribution of PAs. While this limits local applicability, a finer resolution could reveal the specific role of PAs. Future research could investigate the scale dependency of these results. Consequently, our results are intended to illustrate broad-scale patterns rather than serve local-level decision-making (Di Marco et al., 2017; Farooq et al., 2024; Harfoot et al., 2021). In addition, we focused on local predictors within each grid cell and did not account for global externalities, whereby consumption in one country can drive deforestation and biodiversity loss in another through global supply chains (Wiebe and Wilcove, 2025).

Although we included socioeconomic and biophysical variables, these are proxies and cannot capture all pressures directly or fully explain the underlying socioeconomic processes affecting biodiversity loss. Future work could incorporate additional indicators, such as invasive species metrics, to better approximate both direct and indirect drivers of biodiversity loss. The use of static threat maps also limits the detection of temporal change. While PAs appeared to have limited influence in our results, assessing their impact would require temporal threat data, which are currently unavailable (Geldmann et al., 2025; Pulido-Chadid et al., 2025).

Finally, our modeling approach identified the variables most strongly associated with threats to biodiversity and estimated their contribution to the modeled predictions using SHAP values. Thus, we focused on understanding broad patterns, not establishing causation. While SHAP enhances interpretability by showing whether a variable is linked to higher or lower predicted threat probabilities, the analysis remained correlative (Lundberg et al., 2020). The observed associations reflected a co-occurrence rather than a direct causal relationship.

### 4.2 Perspectives

Socioeconomic conditions, followed by LULC, were the strongest predictors of threats across taxa and regions. This reflects a complex interplay of variables, threat types, and taxa, with the influence of human development shaped by income inequality (IPBES, 2019). This highlights broader systems of consumption, governance, and social disparity as key drivers of biodiversity loss, while PAs remain limited and inconsistent in their role.

Our results also highlight that predictors of threats are not uniform across space. The relationships we observed between socioeconomic factors, land cover, and threats were context-dependent, underscoring the need for effective responses to consider contextual and regional differences. Our findings not only highlight the need for improved conservation measures that prioritize areas most in need of protection but also highlight that conservation efforts alone will not be sufficient to halt biodiversity loss. Action must also extend beyond the designation of PAs to address the underlying socioeconomic pressures that degrade ecosystems. To meet the United Nations Sustainable Development Goals (SDGs), particularly SDG 15 (Life on Land) and SDG 12 (Responsible Consumption and Production), integrated approaches that combine conservation and development are necessary.

Halting biodiversity loss will require a shift from isolated conservation actions to a systemic, transformative change that targets both direct and indirect drivers of threats (IPBES, 2025). This means not only strengthening conservation effectiveness but also rethinking our relationship with nature by embedding equity, sustainability, and ecological limits into broader systems of development and governance (Díaz et al., 2019).

