## Supplementary material for "Human development and inequality shape global threat patterns for terrestrial biodiversity"

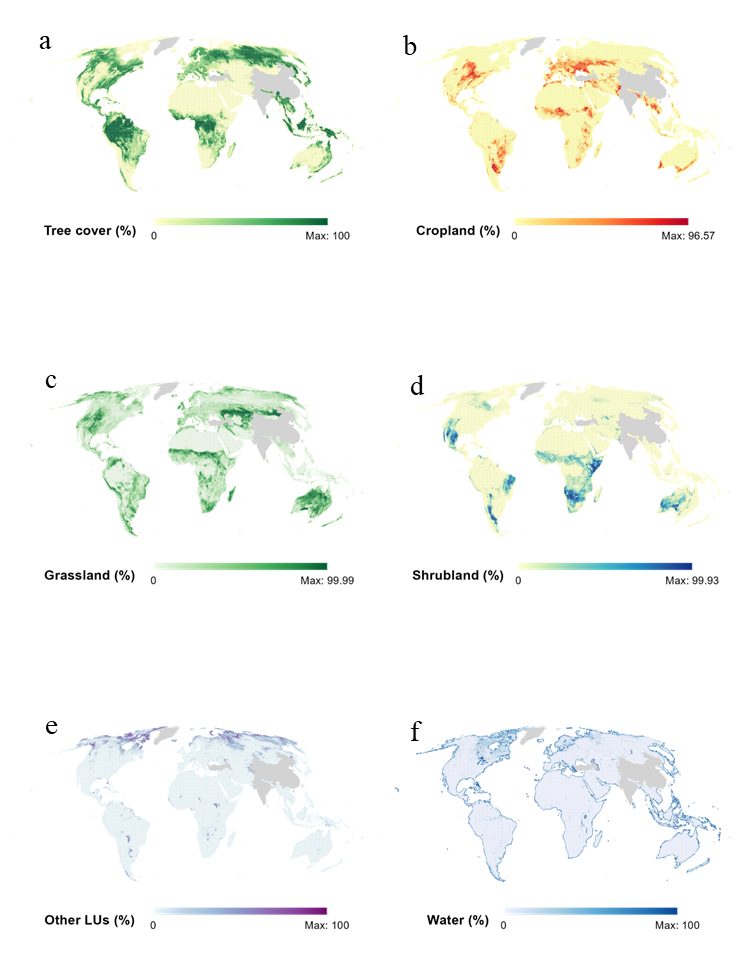

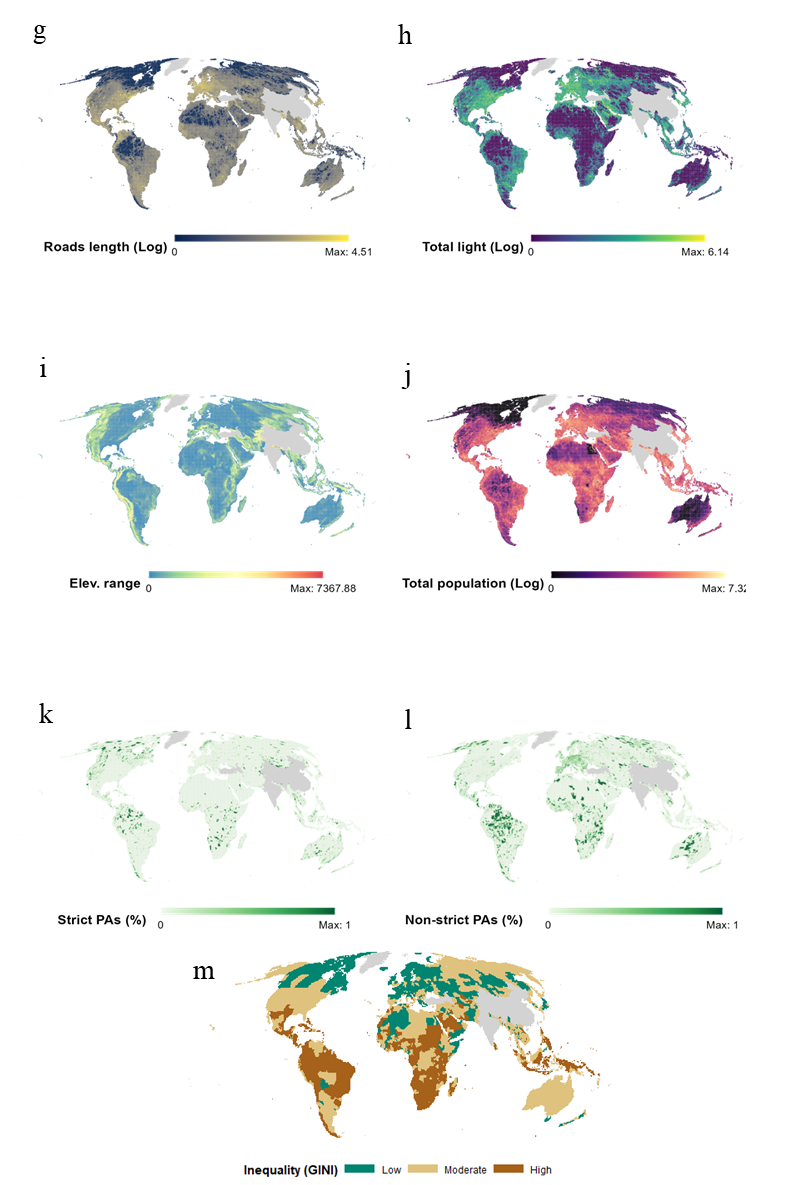

***Figure S1. Transformed variables at 50 × 50 km resolution in Mollweide projection. (a) Percentage tree cover, (b) cropland, (c) grassland, (d) shrubland, (e) other LULC types (including mangroves, ice, and snow), (f) water cover, (g) road length (in km), (h) total nighttime light (logarithmic scale), (i) elevation range, (j) total population count (logarithmic scale), (k) percentage of strict protected area coverage, (l) percentage of non-strict protected area coverage, and (m) GINI index.***

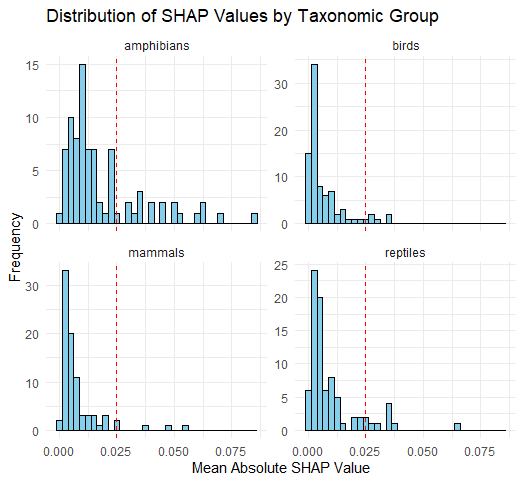

***Figure S2. Distribution of mean absolute SHAP values by taxon for “individual threats models”. Each panel pools mean |SHAP| across the six threat models for that taxon. The red dashed line marks the predefined threshold of 0.025, used to select predictors with high importance and to harmonize comparisons across taxa.***

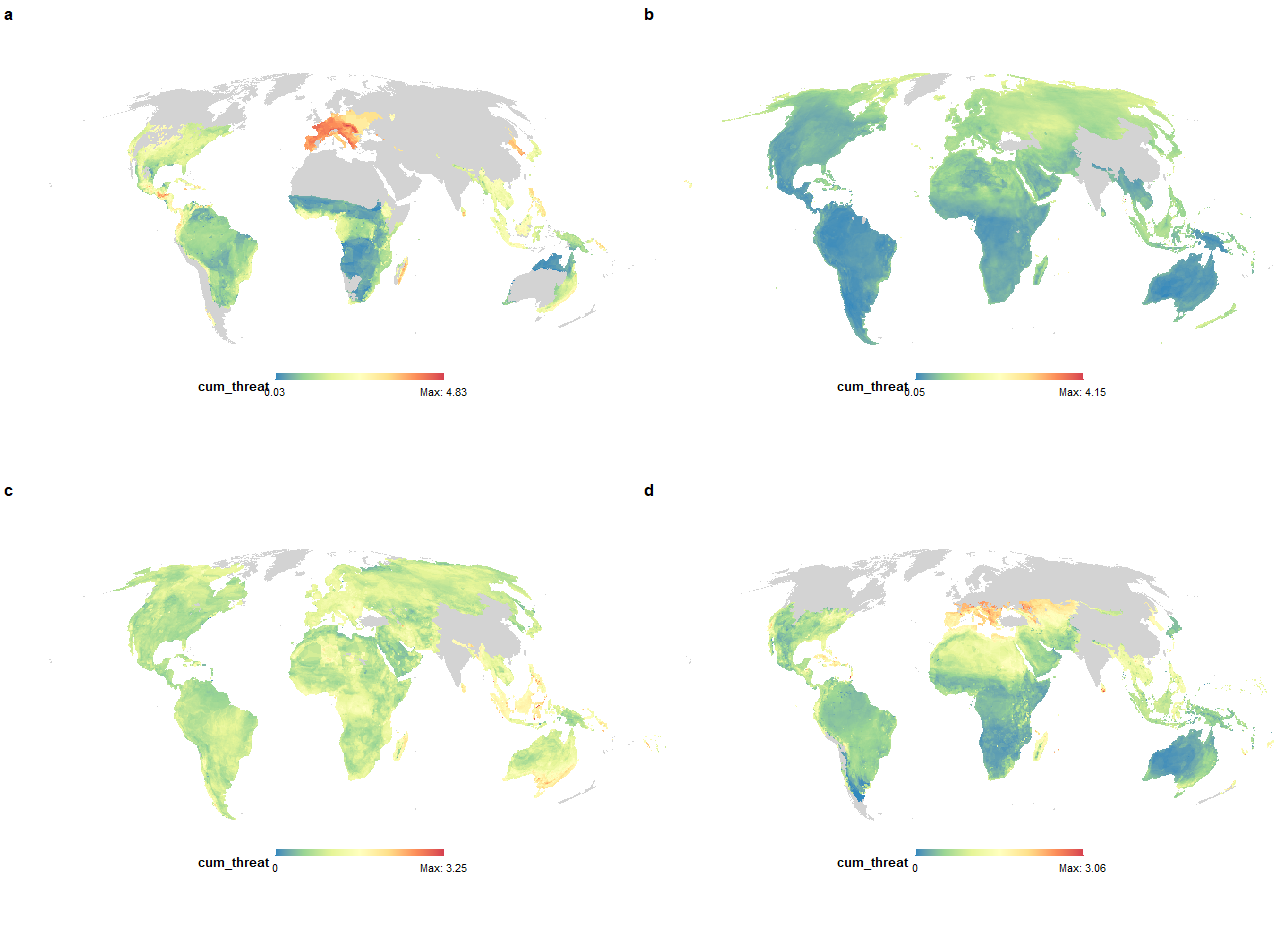

d

b

a

c

***Figure S3. Cumulative threats, represented as the sum of all six threat probabilities per grid cell, for each taxonomic group: (a) amphibians, (b) birds, (c) mammals, and (d) reptiles.***

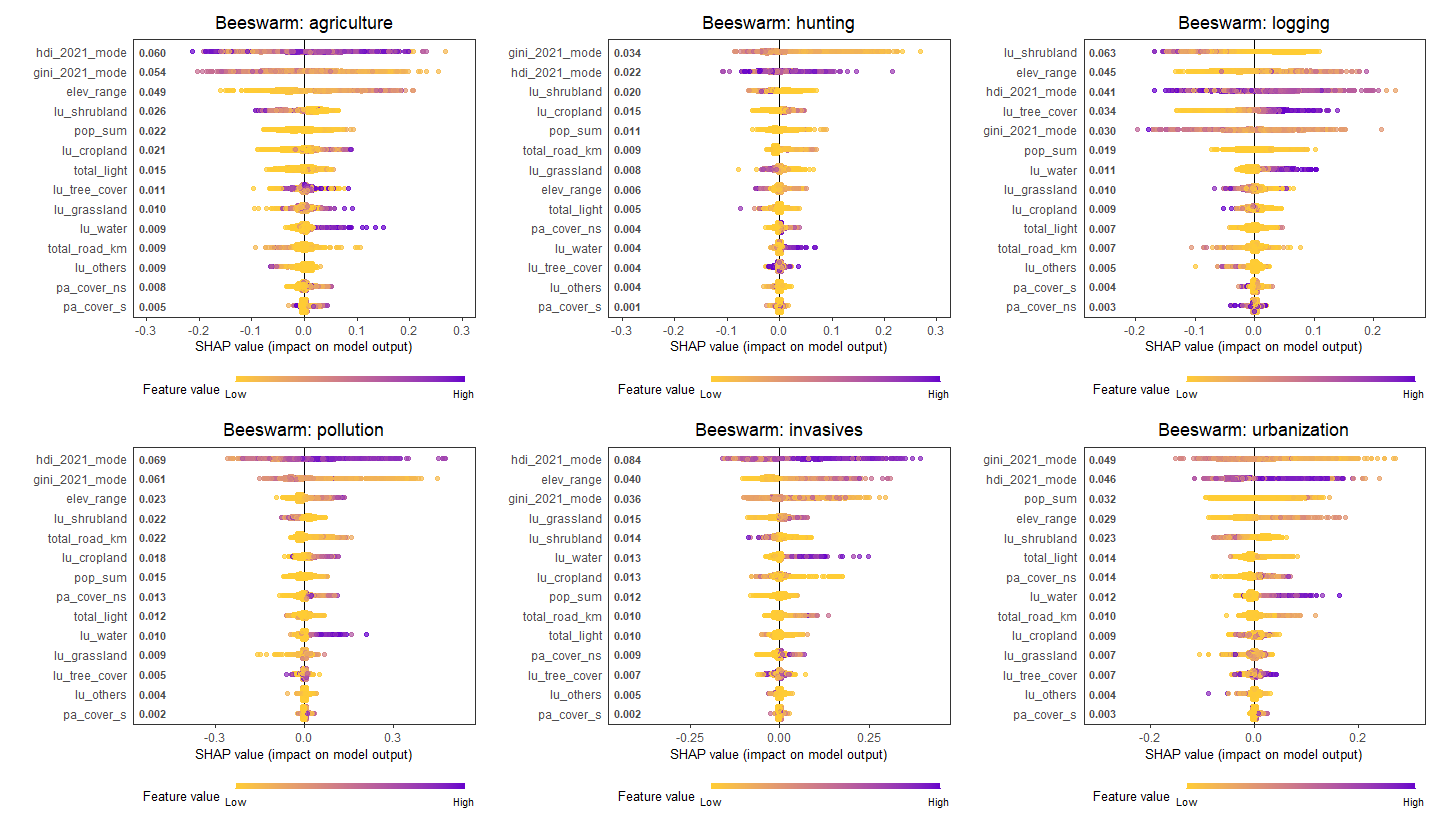

***Figure S4. Beeswarm plot per threat for amphibians. Each plot shows the distribution of SHAP values for all predictors for a given threat, ranked by mean absolute SHAP value. Points represent individual grid cells, with color indicating the actual value of the predictor (purple = low, yellow = high). Positive SHAP values indicate that the predictor increases the predicted probability of the threat, while negative values indicate a decrease, highlighting amphibian-specific patterns in predictor effects across all threats.***

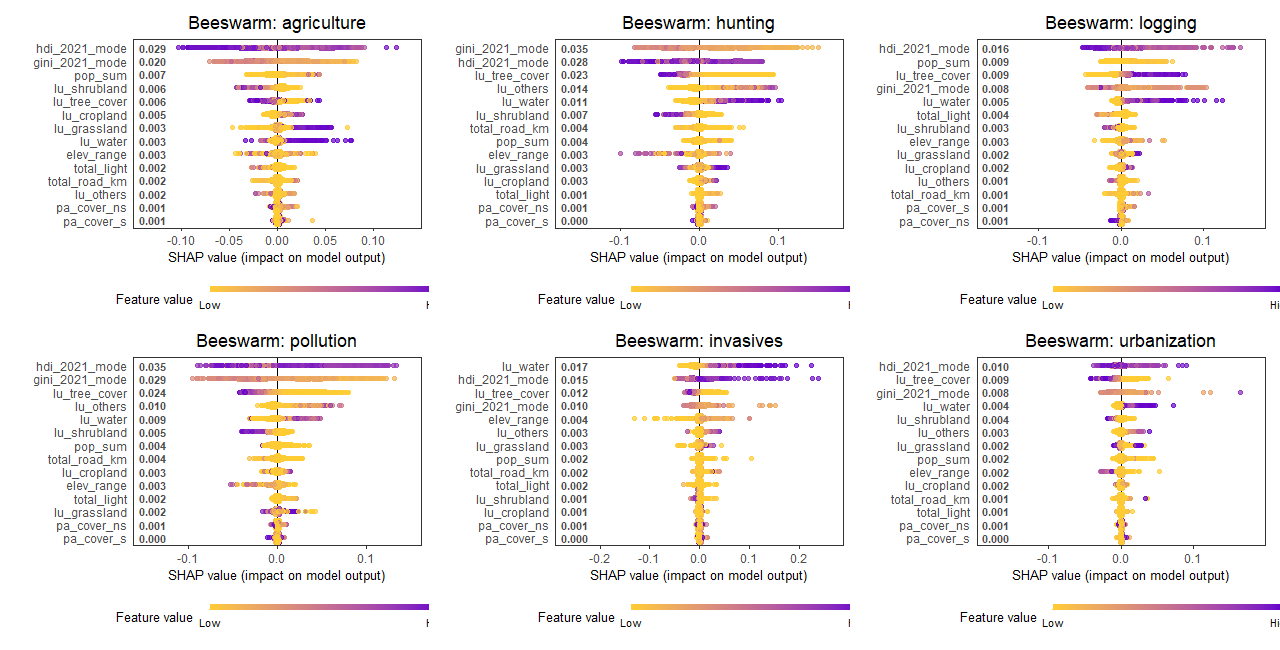

***Figure S5. Beeswarm plot per threat for birds***

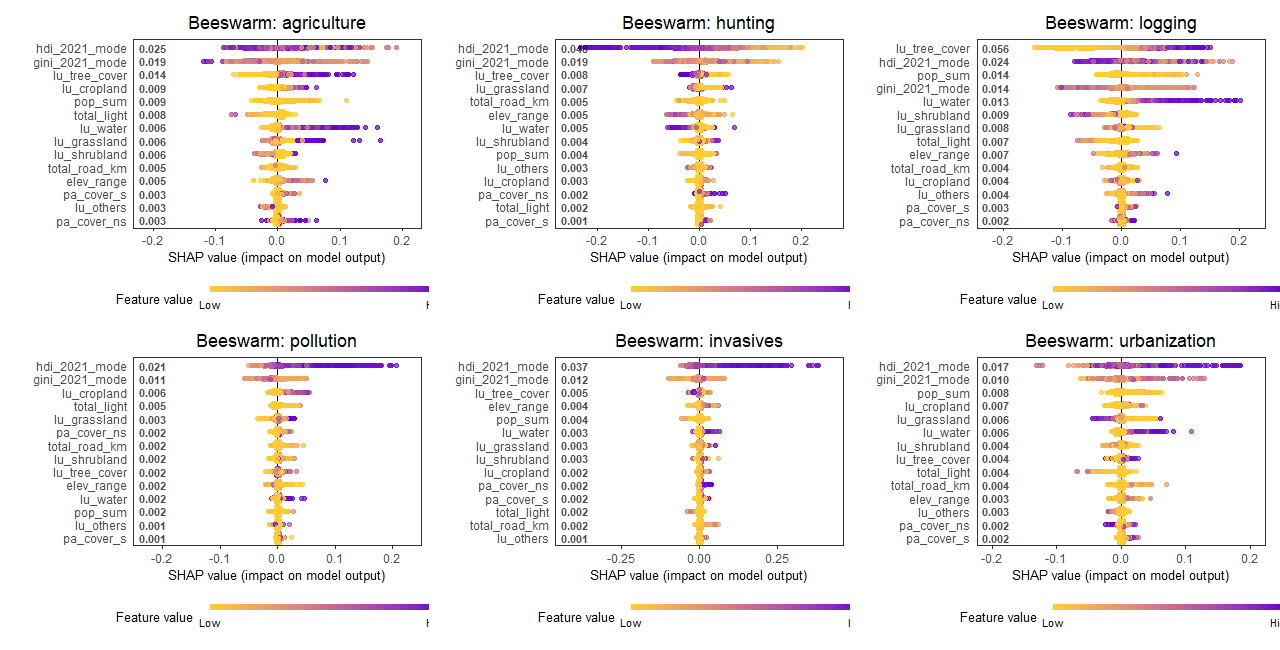

***Figure S6. Beeswarm plot per threat for mammals***

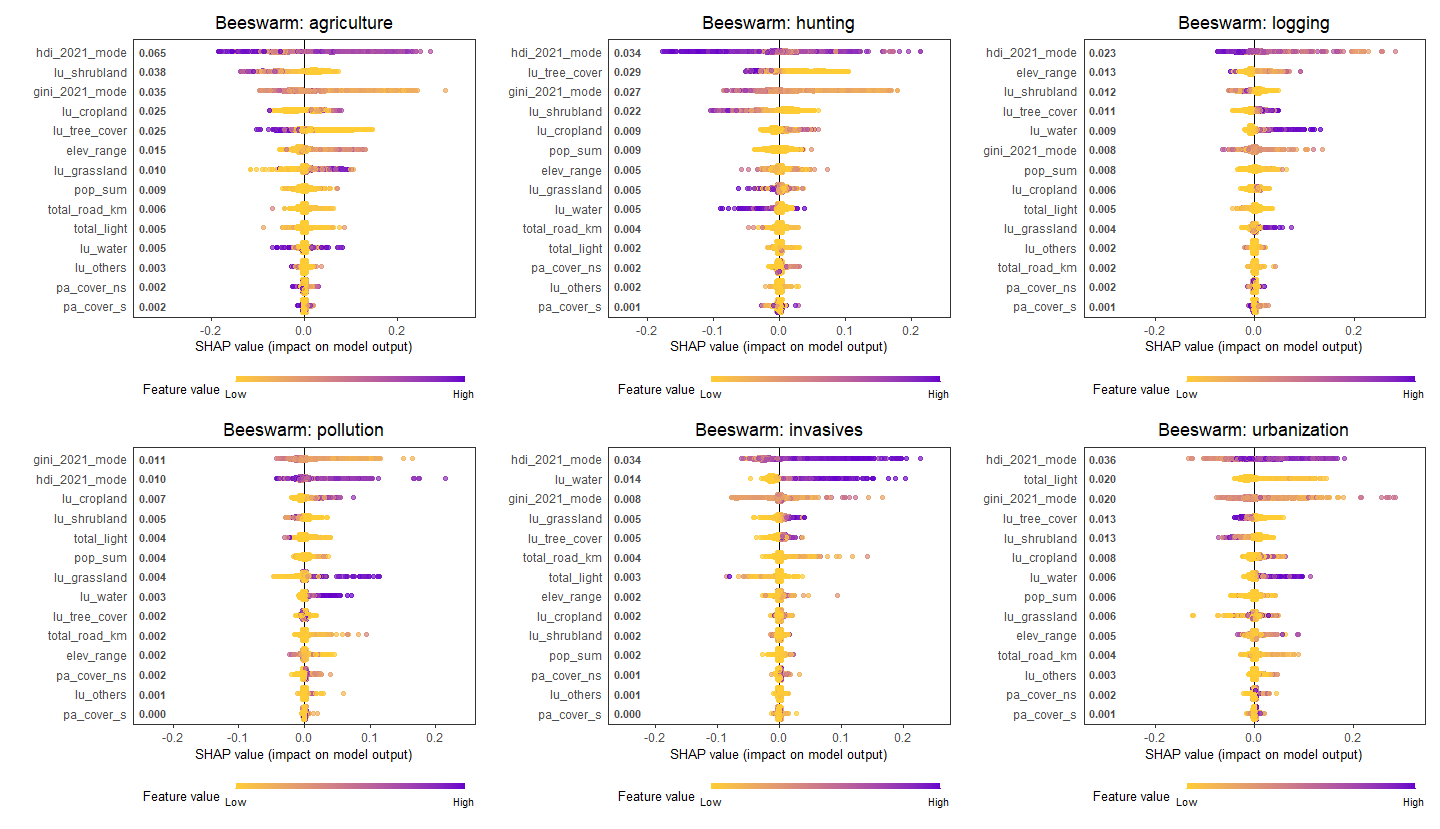

***Figure S7. Beeswarm plot per threat for reptiles***

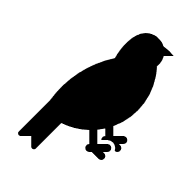

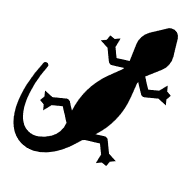

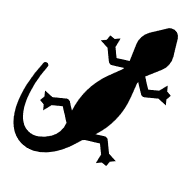

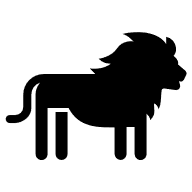

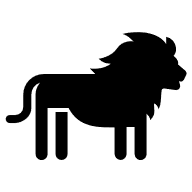

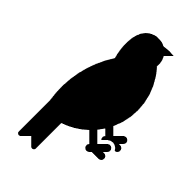

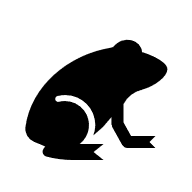

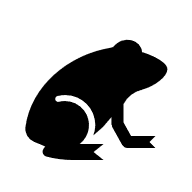

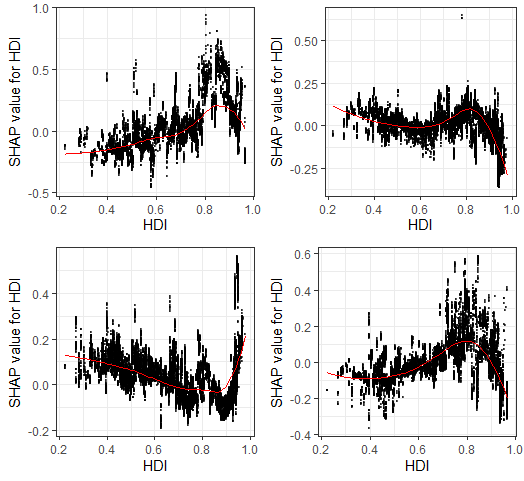

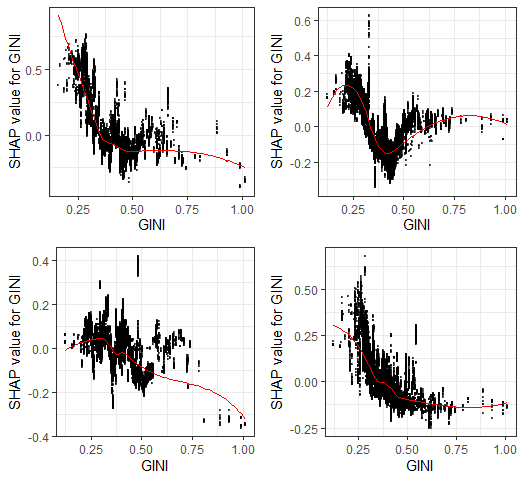

***Figure S8. SHAP dependence plots showing the relationship between the (a) HDI and (b) GINI coefficient and its SHAP value for the cumulative threats model across four taxonomic groups (top left: amphibians, top right: birds, bottom left: mammals, bottom right: reptiles). Each point represents a grid cell, with the x-axis showing the actual HDI/GINI value and the y-axis showing its SHAP value, which reflects the variable’s contribution to the predicted threat probability. Positive SHAP values indicate that higher GINI values contribute to increased predicted threat, while negative SHAP values indicate a contribution to decreased predicted threat. The red line represents a smoothed trend fitted through the SHAP values to highlight the general pattern of the variable’s effect.***

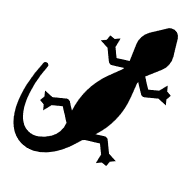

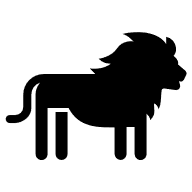

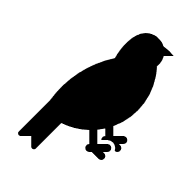

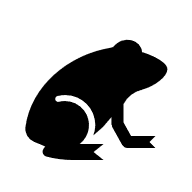

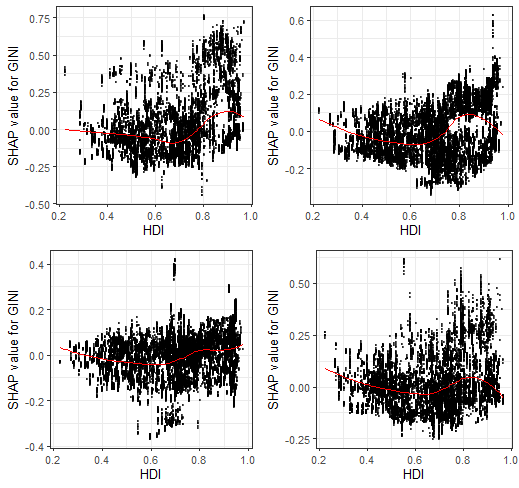

***Figure S9. SHAP dependence plots showing the interaction between Human Development Index (HDI) and the SHAP value for income inequality (GINI) across taxa: a) Amphibians, b) Birds, c) Mammals, and d) Reptiles. Each point represents a grid cell, with the x-axis showing observed HDI values and the y-axis representing the SHAP value of GINI, i.e., the marginal contribution of income inequality to predicted threat probabilities. The red line indicates a smoothed trend (LOESS). Across all taxa, GINI tends to have a stronger positive effect on predicted threats in high-HDI regions, suggesting that income inequality contributes to higher threats in more developed contexts. In contrast, GINI has little or even a slightly negative influence in lower-HDI regions.***

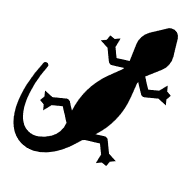

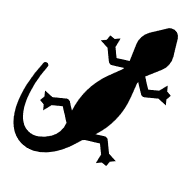

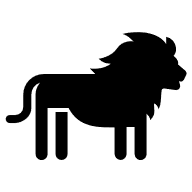

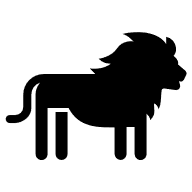

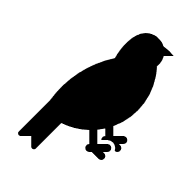

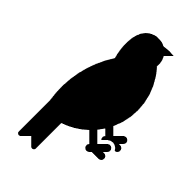

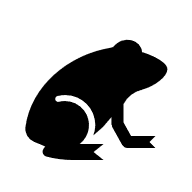

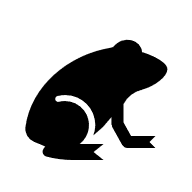

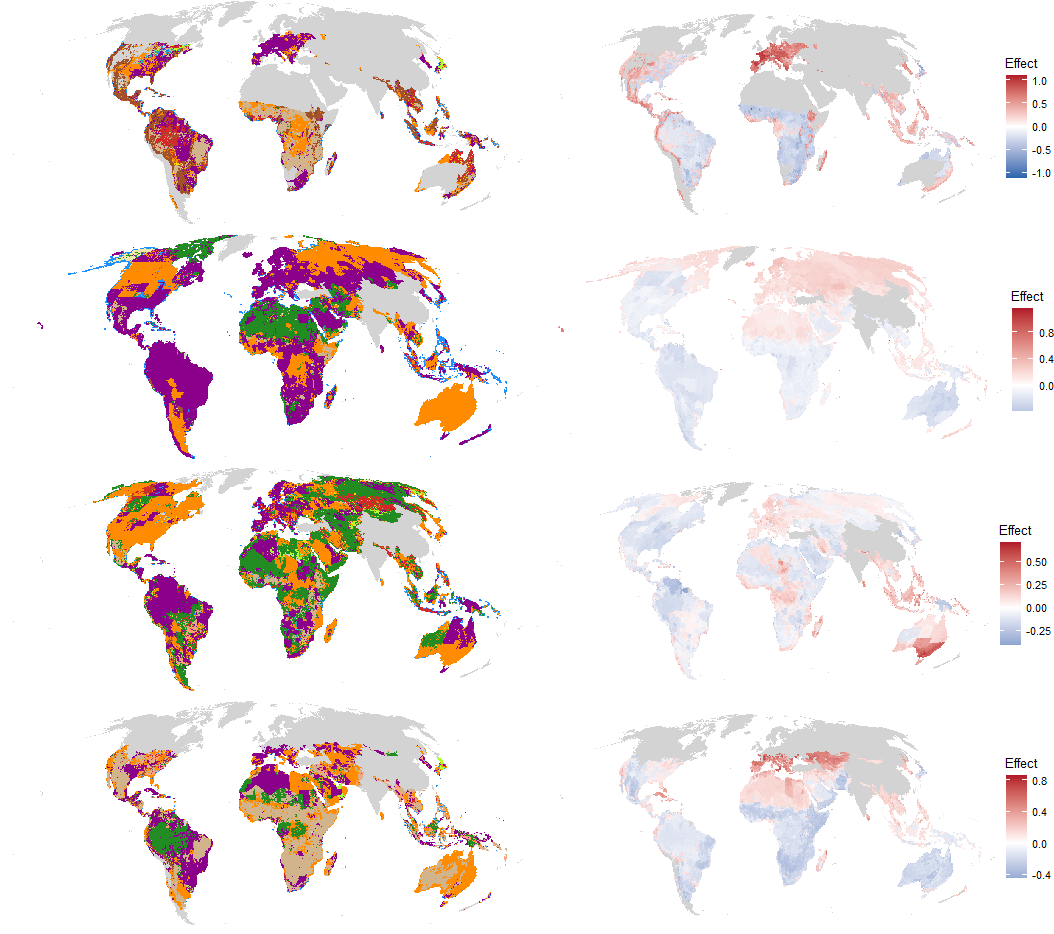

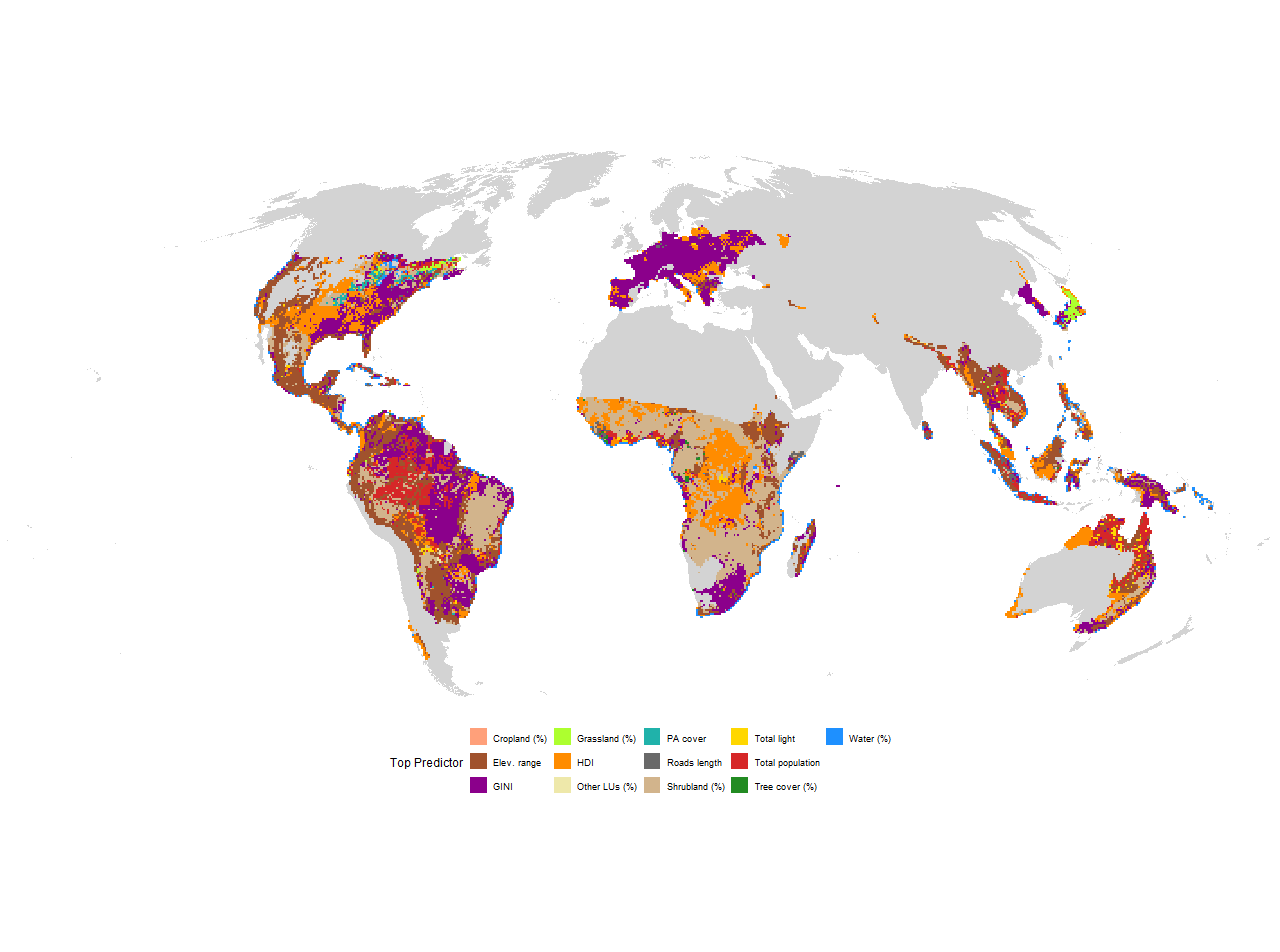

***Figure S10. Variables with the highest relative importance according to SHAP values for the cumulative threats model. Maps on the left show the spatial distribution of the top predictor. Maps on the right show the mean SHAP value for that predictor across grid cells, indicating its effect on the predicted threat: red areas are locations where the predictor contributes to higher predicted threat probability, while blue areas are locations where it contributes to lower predicted threat probability.***

### Model performance

Among the individual threats, the highest R² values were for birds, with pollution and hunting models reaching 0.94 and 0.91, respectively. In contrast, invasive species predictions for reptiles had the lowest R² (0.66), suggesting higher variability or weaker associations with the selected predictors. Cumulative threats models performed comparably or better than the individual threat models, particularly for amphibians (R² = 0.88) and birds (R² = 0.89). Their RMSEs remained low relative to the scale of the cumulative threat scores, indicating low prediction error in proportion to the maximum values threat values (Table S1).

***Table S1. Model performance for individual and cumulative threats per taxon. Performance metrics are reported as RMSE and R² values. Models were selected based on the best performance after hyperparameter tuning.***

| Taxon | Step | Target | Max. threat value | RMSE | R^2^ |
| --- | --- | --- | --- | --- | --- |
| Amphibians | 1. Individual threats | Agriculture | 1 | 0.11 | 0.81 |
|  | 1. Individual threats | Hunting | 0.9 | 0.06 | 0.82 |
|  | 1. Individual threats | Invasives | 1 | 0.09 | 0.85 |
|  | 1. Individual threats | Logging | 1 | 0.1 | 0.78 |
|  | 1. Individual threats | Pollution | 1 | 0.09 | 0.89 |
|  | 1. Individual threats | Urbanization | 0.99 | 0.1 | 0.79 |
|  | 2. Cumulative | Cumulative threats | 4.83 | 0.35 | 0.88 |
| Birds | 1. Individual threats | Agriculture | 0.94 | 0.03 | 0.84 |
|  | 1. Individual threats | Hunting | 0.82 | 0.03 | 0.91 |
|  | 1. Individual threats | Invasives | 0.96 | 0.03 | 0.81 |
|  | 1. Individual threats | Logging | 0.85 | 0.03 | 0.78 |
|  | 1. Individual threats | Pollution | 0.74 | 0.03 | 0.94 |
|  | 1. Individual threats | Urbanization | 0.86 | 0.02 | 0.74 |
|  | 2. Cumulative | Cumulative threats | 4.15 | 0.11 | 0.89 |
| Mammals | 1. Individual threats | Agriculture | 0.95 | 0.06 | 0.71 |
|  | 1. Individual threats | Hunting | 0.91 | 0.05 | 0.82 |
|  | 1. Individual threats | Invasives | 0.91 | 0.04 | 0.83 |
|  | 1. Individual threats | Logging | 0.92 | 0.05 | 0.83 |
|  | 1. Individual threats | Pollution | 0.51 | 0.02 | 0.84 |
|  | 1. Individual threats | Urbanization | 0.79 | 0.04 | 0.69 |
|  | 2. Cumulative | Cumulative threats | 3.25 | 0.16 | 0.71 |
| Reptiles | 1. Individual threats | Agriculture | 0.98 | 0.08 | 0.84 |
|  | 1. Individual threats | Hunting | 0.88 | 0.05 | 0.84 |
|  | 1. Individual threats | Invasives | 1 | 0.04 | 0.66 |
|  | 1. Individual threats | Logging | 0.88 | 0.05 | 0.72 |
|  | 1. Individual threats | Pollution | 0.75 | 0.03 | 0.77 |
|  | 1. Individual threats | Urbanization | 0.9 | 0.06 | 0.78 |
|  | 2. Cumulative | Cumulative threats | 3.06 | 0.19 | 0.85 |
